# Structural dissection of the first events following membrane binding of the islet amyloid polypeptide

**DOI:** 10.1101/2021.12.14.472560

**Authors:** Lucie Khemtemourian, Hebah Fatafta, Benoit Davion, Sophie Lecomte, Sabine Castano, Birgit Strodel

**Affiliations:** Université de Bordeaux, CNRS, Bordeaux INP, CBMN, UMR 5248, F-33600 Pessac, France; Institute of Biological Information Processing: Structural Biochemistry, Forschungszentrum Jülich, 52428 Jülich, Germany; JuStruct, Jülich Center for Structural Biology, Forschungszentrum Jülich, 52428 Jülich, Germany; Institute of Theoretical and Computational Chemistry, Heinrich Heine University Düsseldorf, 40225 Düsseldorf, Germany

**Keywords:** islet amyloid polypeptide, type 2 diabetes mellitus, amylin, amyloid aggregation, peptide-membrane interactions

## Abstract

Amyloid forming proteins are involved in many pathologies and often belong to the class of intrinsically disordered proteins. One of these proteins is the islet amyloid polypeptide (IAPP), which is the main constituent of the amyloid fibrils found in the pancreas of type 2 diabetes patients. The molecular mechanism of IAPP-induced cell death is not yet understood, however it is known that cell membrane plays a dual role, being a catalyst of IAPP aggregation and the target of IAPP toxicity. Using FTIR spectroscopy, transmission electron microscopy, and molecular dynamics simulations we investigate the very first molecular steps following IAPP binding to a lipid membrane. In particular, we assess the combined effects of the charge state of amino-acid residue 18 and the IAPP-membrane interactions on the structures of monomeric and aggregated IAPP. Both our experiments and simulations reveal distinct IAPP-membrane interaction modes for the various IAPP variants. Membrane binding causes IAPP to fold into an amphipathic helix, which in the case of H18K- and H18R-IAPP can easily insert into the membrane. For all IAPP variants but H18E-IAPP, the membrane-bound *α*-helical structure is an intermediate on the way to IAPP amyloid aggregation, while H18E-IAPP remains in a stable helical conformation. The fibrillar aggregates of wild-type IAPP and H18K-IAPP are dominated by an antiparallel *β*-sheet conformation, while H18R- and H18A-IAPP exhibit both antiparallel and parallel *β*-sheets as well as amorphous aggregates. In summary, our results emphasize the importance of residue 18 for the structure and membrane interaction of IAPP. This residue is thus a good target for destabilizing amyloid fibrils of IAPP and inhibit its toxic actions by possible therapeutic molecules.

## 1 INTRODUCTION

The formation of amyloid fibrils is involved in various human diseases, such as Alzheimer’s disease, Parkinson’s disease, or type 2 diabetes mellitus (T2DM). Amyloid forming proteins are often intrinsically disordered proteins (IDPs) or are proteins that contain one or more intrinsically disordered regions. The structure of those amyloid fibrils are very heterogeneous but they are all composed of arrays of cross *β*-sheets Selkoe (2004); Knowles et al. (2014)

The human islet amyloid polypeptide (hIAPP), also known as amylin, is a 37-amino acid peptide hormone that is the main constituent of the islet amyloid mainly found in the pancreatic islets of T2DM patients, but also in many organs including the brain, the heart, and the kidney Westermark et al. (1987); Cooper et al. (1988); de Koning et al. (1995); Despa et al. (2012); Srodulski et al. (2014) hIAPP is produced and secreted together with insulin by the pancreatic *β*-cells, and it plays a role in the control of glucose homeostasis and satiety by acting on the liver, gut, brain and pancreas Lutz (2010); Westermark et al. (2011). Under normal conditions, monomeric hIAPP lacks a well-defined structure as typical for an IDP and mainly adopts a random coil conformation. However, in T2DM patients, hIAPP starts to aggregate into amyloid fibrils and the formation of these amyloid aggregates has been associated with the dysfunction and death of *β*-cells Opie (1901); Höppener et al. (2000).

While the toxic activity of hIAPP is still not completely understood, a link between hIAPP fibril formation at the membrane interface and hIAPP-induced cell death was observed, highlighting the relevance of the membrane. A few putative mechanisms of cell membrane-disruption by hIAPP have been described and have been the subject of several studies Mirzabekov et al. (1996); Janson et al. (1999); Hebda and Miranker (2009); Engel et al. (2008); Martel et al. (2016) It has been suggested that the amyloid fibrils are not the primary toxic species, but the oligomers formed by hIAPP are thought to be cytotoxic, either by forming membrane channels or by inducing bilayer disorder Mirzabekov et al. (1996); Quist et al. (2005); Kayed et al. (2004) In agreement with these studies, molecular dynamics (MD) simulations demonstrated that membrane permeability was induced by oligomeric hIAPP Poojari et al. (2013). Further experimental studies have indicated that the formation of hIAPP fibrils at the membrane causes membrane disruption by forcing the curvature of the bilayer to unfavorable angles or by the uptake of lipids by the fibrils Engel et al. (2008); Sparr et al. (2004). Even if the mechanism is not yet fully understood, altogether these studies revealed the importance of the membrane in hIAPP-induced cell death.

Along with these results, it has been recognized that the various amino acids of hIAPP are crucial in hIAPP fibril formation and in hIAPP-membrane disruption. The N-terminal part residues are mainly responsible for membrane binding, the middle core drives amyloid fibril formation, while the C-terminal residues are also involved in amyloid fibril formation yet to a lesser extent Skeby et al. (2016); Engel et al. (2006); Brender et al. (2008a,b). The sequence of IAPP is highly conserved across different species Caillon et al. (2016); Cao et al. (2013), however key differences, that play important roles in modulating the propensity of the peptide to aggregate, have been identified. The non-amyloidogenic and non-toxic mouse IAPP differs from hIAPP by six residues out of 37; Interestingly, five of the six residues are located in the amyloid-prone region 20–29 and mice do not develop T2DM. For that reason, it is essential to explore the sequence-structure relationship. While the region 20–29 is of relevance, it is not the sole region governing IAPP fibril formation, since proline mutations at position 14, 15, 16 or 17 can also induce a loss of fibril formation Abedini and Raleigh (2006); Fox et al. (2010); Tu and Raleigh (2012). Recent studies on residue 18, that is highly variable among species Caillon et al. (2016), indicate that this residue is important in modulating i) IAPP fibril formation in solution and in the presence of membranes Khemtemourian et al. (2017); Hoffmann et al. (2018), ii) membrane interaction and damage Hoffmann et al. (2018), iii) cell toxicity Khemtemourian et al. (2017), and iv) hIAPP-zinc and hIAPP-insulin affinity Khemtemourian et al. (2021).

The characterization of the aggregation pathways and of the structure at a molecular and an atomic level at the membrane is thus a key step to understand hIAPP cellular toxicity and its role in disease states. While structure of hIAPP in solution was extensively studied Goldsbury et al. (2000); Camargo et al. (2017); Williamson and Miranker (2007); Wiltzius et al. (2009), only a few studies were performed in a membrane environment, which mainly using spectroscopic techniques such as circular dichroism (CD) or nuclear magnetic resonance (NMR) spectroscopy Jayasinghe and Langen (2005); Milardi et al. (2021); Caillon et al. (2013); Nanga et al. (2011); Patil et al. (2009); Camargo et al. (2017). Both CD and NMR techniques provide information on structural averages of the conformational ensemble. A complicating aspect for NMR spectroscopy of hIAPP in the presence of lipid bilayers is the fast aggregation speed of hIAPP. To overcome this challenge, approaches have been developed to hamper the fibrillation process, such as the use of low temperatures and/or detergent micelles that stabilize the monomeric form of hIAPP Jayasinghe and Langen (2005); Caillon et al. (2013); Nanga et al. (2011); Patil et al. (2009); Camargo et al. (2017). Here, we address this problem by employing a combination of two techniques that offer the possibility of obtaining time-resolved structural information of hIAPP in a membrane environment, namely Fourier-transform infrared spectroscopy (FTIR) spectroscopy and MD simulations. This allows us to provide structural information for both monomeric hIAPP as well as the first aggregations of hIAPP at the membrane. Previous simulation studies examined the membrane interactions of monomeric and oligomeric hIAPP Dignon et al. (2017a); Dong et al. (2018); Martel et al. (2016); Press-Sandler and Miller (2018); Qian et al. (2018); Qiao et al. (2019). The results from these simulations indicate that wild-type hIAPP interacts with the membrane by forming interactions between the anionic lipids of the membrane and the N-terminal part of hIAPP, which is in agreement with experimental data Skeby et al. (2016); Engel et al. (2006). Stabilization of the *α*-helical state following the binding to a membrane was also observed in both experimental and simulation studies Caillon et al. (2013); Dignon et al. (2017a) FTIR spectroscopy has been previously used to provide insights into the membrane-bound monomeric and fibril structures of hIAPP Mishra et al. (2008); Mishra and Winter (2008); Radovan et al. (2008). The studies indicated that a transition from unordered structures to *β*-sheet structures occurs on a time scale characteristic for amyloid fibril formation.

The purpose of this study is to obtain structural information on native hIAPP at the membrane interface and to determine the role of histidine 18 in hIAPP-membrane interactions. All the experiments were performed with wild-type hIAPP and four mutated peptides where histidine 18 has been replaced by arginine (H18R-IAPP), lysine (H18K-IAPP), glutamic acid (H18E-IAPP), and alanine (H18A-IAPP) to achieve variations in charge, shape, volume, and hydrophobicity. To evaluate the interaction of hIAPP and the mutated peptides with the membrane, we worked with a 1,2-dioleoyl-sn-glycero-3-phosphocholine/1,2-dioleoyl-sn-glycero-3-phospho-L-serine (DOPC/DOPS) lipid mixture (ratio 7:3) to mimic the eukaryotic *β*-cell membranes. These cells contain typically between 1–10% of negatively charged lipids; however in the case of T2DM, the high concentration of glucose increasex the amount of negatively charged lipids up to 30% Rustenbeck et al. (1994). We performed attenuated total reflection (ATR) FTIR spectroscopy at different incubation times to apprehend the initial structure of the peptides at the membrane and the evolution of structural changes. The putative perturbation of the lipid membranes after addition of the peptides was also investigated. We observed differences for the wild-type and the mutated peptides not only in the initial structures but also in the variation of secondary structure in time, highlighting the role of the residue histidine 18 in the membrane interactions of hIAPP and in the process of fibril formation. The ATR-FTIR results are augmented on either side of the length and time scales, by MD simulations to provide mechanistic insight in the structural transitions and peptide-membrane interactions and by transmission electron microscopy (TEM) to obtain images of the final fibrils.

## 2 MATERIALS AND METHODS

### 2.1 Sample preparation

Peptide solutions were prepared as described previously Hoffmann et al. (2018); Khemtemourian et al. (2017). Briefly, stock solutions were obtained by dissolving the peptide powder at a concentration of 1 mM in hexafluoroisopropanol (HFIP) and by letting them incubate for an hour. HFIP was then evaporated under a stream of dry N_2_ and further dried by vacuum in a desiccator for at least 30 min. The resulting peptide film was then rehydrated with 100 *µ*L of buffer and 2 *µ*L of a CaCl_2_ solution.

### 2.2 Preparation of phospholipid vesicles

DOPC and DOPS lipids were purchased from Avanti Polar Lipids. Lipid powders were dissolved in chloroform and mixed at the desired ratio. The solvent was evaporated under a stream of dry nitrogen and further dried under high vacuum in a desiccator for at least 30 min. Lipid films were then rehydrated for 1 h with a buffer of 10 mM Tris, 100 mM NaCl, pH 7.4 in 100% D_2_O, obtaining large, multilamellar vesicles (LMVs). Small, unilamellar vesicles (SUVs) were then prepared from the LMVs by tip sonication. The SUVs were burst onto a germanium ATR crystal to form a single bilayer which is controlled by the measurement of the absolute IR intensity. Large, unilamellar vesicles (LUVs) were prepared using the same buffer conditions as for the LMVs, but containing 100% H_2_O, which was subjected to 10 freeze-thaw cycles with alternating temperatures of about −190^°^C and 50^°^C. The lipid suspension was subsequently extruded 19 times through a mini-extruder (Avanti Polar Lipids) equipped with a 200 nm polycarbonate membrane. The phospholipid content of both lipid stock solutions and vesicles was determined as inorganic phosphate according to Rouser et al. Rouser et al. (1970)

### 2.3 ATR-FTIR spectroscopy

ATR-FTIR spectra were recorded on a Nicolet 6700 spectrometer Thermo Scientific equipped with an MCT detector cooled at 77 K. A Ge-crystal was used as internal reflection unit. Since ATR-FTIR spectroscopy is sensitive to the orientation of the structures Goormaghtigh et al. (1990, 1994, 1999), spectra were recorded with parallel (p) and perpendicular (s) polarizations of the incident light with respect to the ATR plate. 200 scans were recorded at a resolution of 8 cm^−1^. All the orientation information is then contained in the dichroic ratio *R*_ATR_ = *A*_p_*/A*_s_, where *A*_p_ and *A*_s_ represent the absorbance underlying the band at p and s, respectively, polarization of the incident light. After subtraction of a spectrum of the lipid membrane with the buffer and subtraction of noise from water, the spectra were baseline-corrected between 1700 and 1600 cm^−1^ corresponding to the amide I band area. Finally, a smoothing has been applied. To derive the secondary structure from the bands, the spectra were analyzed with an algorithm based on a second-derivative function and a self-deconvolution procedure (GRAMS and OMNIC software, Thermo Fisher Scientific) to determine the number and wavenumber of the individual bands within the spectral range of the amide I band.

### 2.4 Transmission electron microscopy

TEM was performed at the “Institut de Biologie Paris Seine” (IBPS, Sorbonne Université, Paris, France). Peptides and LUVs were incubated for 2 days at room temperature. Aliquots (20 *µ*L) were adsorbed onto a glow-discharged carbon coated 200 mesh copper grid for 2 minutes and then negatively stained with saturated uranyl acetate for 45 seconds. Grids were examined using a ZEISS 912 Omega electron microscope operating at 80 kV.

### 2.5 Computational methods

#### 2.5.1 Setup of the simulated systems

The modeled systems are composed of the full-length (37 residues) hIAPP monomer (either wild-type or mutated at residue 18) and a DOPC/DOPS lipid bilayer in a 7:3 ratio mimicking the lipid composition of the experiments. As initial peptide structure, the most populated conformation from a preceding 1 *µ*s simulation of wild-type hIAPP as a monomer and with a disulfide bond between C2 and C7 in the aqueous phase was used. The mutated peptides were generated from this structure by replacing the neutral H18 residue (protonated only at N*o*) by its positively charged counterpart (denoted by H18+), the neutral residue alanine, the negatively charged glutamate, or the positively charged lysine or arginine using the CHARMM-GUI interface (Lee et al., 2016). These peptides will be referred to as hIAPP, hIAPP(H18+), H18A-IAPP, H18E-IAPP, H18K-IAPP, and H18R-IAPP, respectively. CHARMM-GUI was also used to set up and equilibrate the DOPC/DOPS lipid bilayer as a symmetric membrane composed of 88 DOPC and 40 DOPS lipid molecules. The peptides were placed above the lipid bilayer (one peptide per simulation) at a distance of ≈3 nm from the bilayer surface. Each system was then solvated with water using the TIP3P model and NaCl was added at physiological concentration of 150 mM, while also neutralizing the system. The total number of atoms *N* in each system was ≈54,000 atoms and the simulation box size was about 6.5 × 6.5 × 12.0 nm^3^.

#### 2.5.2 MD simulation conditions

The MD simulations were carried out using the GROMACS 2018.2 simulation package (Abraham et al., 2015), along with CHARMM36 (Klauda et al., 2010) as force field for the lipids and CHARMM36m (Huang et al., 2017) for the IAPP peptides. Each system was first energy minimized using the steepest descent algorithm to remove initial atom clashes that may have resulted during the setup. This was followed by an equilibration using MD simulations under *NVT* conditions, where the reference temperature *T* of 302 K (which was chosen to be close to the temperatures used in the experiments) was regulated with a velocity-rescale thermostat (Bussi et al., 2007). Then, the system was equilibrated under *NpT* conditions to obtain a pressure *p* of 1.0 bar, which was realized by regulating the pressure using a semi-isotropic Parrinello-Rahman pressure coupling scheme (Berendsen et al., 1984). The particle mesh Ewald (PME) method was used to calculate the electrostatic interactions in combination with periodic boundary conditions set in all directions. The electrostatic interactions in real space as well as the van der Waals interactions were cut at 1.2 nm. All bonds were constrained using the LINCS algorithm. For each of the six systems the MD simulations were run in triplicate and for 1 *µ*s per simulation (i.e., 3 × 1 *µ*s per system).

#### 2.5.3 Analysis of the MD simulations

All analysis programs mentioned are available via the GROMACS 2018.2 program package (Abraham et al., 2015). Only the MD snapshots were IAPP is within 0.5 nm of the bilayer were included in the analysis of the system in question. The peptide-lipid interactions were then determined by calculating the interaction energy between each IAPP residue and the DOPC and DOPS lipids, respectively, using ‘gmx energy’. The ‘gmx mindist’ program was employed to determine the number of contacts between each IAPP residue and DOPC/DOPS. A contact was recorded when the distance between any two non-hydrogen atoms from a residue and a lipid was within 0.5 nm. The hydrogen bond propensity was determined as the ratio of the number of MD snapshots where one or more hydrogen bonds were formed between peptide and lipid and the total number of MD snapshots per system. The secondary structure of the peptides was determined using the ‘define secondary structure program’ (DSSP) invoked via the GROMACS tool ‘do dssp’. To facilitate a clear representation, the data of similar secondary structures are grouped together: *β*-strand and *β*-bridge are combined as *β*-sheet, *β*-turn and bend as turn, and helix includes *α*-, *π*- and 3_10_-helices.

## 3 RESULTS

### 3.1 hIAPP and the mutated peptides are initially unstructured at the membrane interface

We first investigated, using ATR-FTIR spectroscopy, the structural behavior of wild-type and mutant hIAPP when interacting with a (supported) lipid bilayer composed of DOPC/DOPS (7:3). These phospholipids represent the most abundant zwitterionic phospholipid species (PC) and the dominant negatively charged phospholipid species (PS) in eukaryotic cells, and the 7:3 ratio is similar to the one of zwitterionic lipids to negatively charged lipids of the membrane of pancreatic islet cells Rustenbeck et al. (1994). We performed polarized ATR-FTIR experiments in order to analyze the initial structures of peptides at the membrane and to determine if the mutation at residue 18 could induce some structural changes. Figure 1 shows the ATR-FTIR spectra in the amide I region of hIAPP and the mutated peptides interacting with DOPC/DOPS bilayers. Based on the amide I band analysis, hIAPP and H18K-IAPP exhibit two peaks at around 1643 ± 1 cm^−1^ and 1623 ± 1 cm^−1^ assigned to random coil and *β*-sheet structure, respectively. The mutated peptides H18R-IAPP, H18E-IAPP and H18A-IAPP display predominantly an amide I band at around 1643 ± 1 cm^−1^ that can be attributed to unordered secondary structures. The secondary structure content of the peptides bound to the DOPC/DOPS bilayers has then been evaluated from the analysis of the amide I band shape and curve fitting (Table 1). The bands at 1686 ± 1 cm^−1^, 1632 ± 1 cm1^−1^, 1624 ± 1 cm^−1^, and 1615 ± 1 cm^−1^ were assigned to *β*-sheets (parallel and antiparallel), the band at 1654 ± 1 cm^−1^ to *α*-helices, the band at 1643 ± 1 cm^−1^ to random structures, and the one at 1674 ± 1 cm^−1^ to *β*-turns. The results show that hIAPP is mainly unstructured (40%) with a contribution of *β*-sheets (31%), which is in agreement with previous studies Khemtémourian et al. (2010); Seeliger et al. (2012). The peptides H18R-IAPP and H18K-IAPP adopt unstructured conformation with about the same probability as hIAPP (41% and 37%, respectively) but have different amounts of *β*-sheets (27% and 36%). The initial structure of H18E-IAPP differs substantially from the wild-type peptide with less *β*-sheet content and more random coil conformation. The peptide H18A-IAPP has the highest content of *α*-helical structure and the lowest amount of random coil, which is likely due to the inherent preference of alanine to adopt a helical conformation; in fact, alanine is regarded as the most stabilizing residue in helices. Such change in the initial structure may modify the kinetics of fibril formation as shown previously Hoffmann et al. (2018). Overall, the data indicate that at the membrane interface, hIAPP is initially largely unstructured, but depending on the kind of mutation at residue position 18, the peptides also adopt *β*-sheet and *α*-helical structures to different extents. In order to determine if these mutations do also influence the kinetics of structural changes, the same kind of experiments were carried out during the course of a few hours.

**Table 1.**
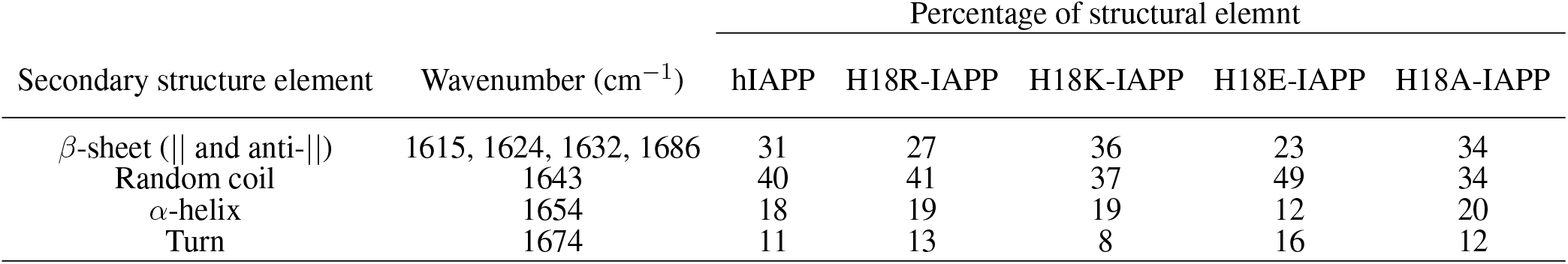
Secondary structure content derived from ATR-FTIR spectra of hIAPP and its mutants interacting with a DOPC/DOPS membranes.

**Figure 1.**
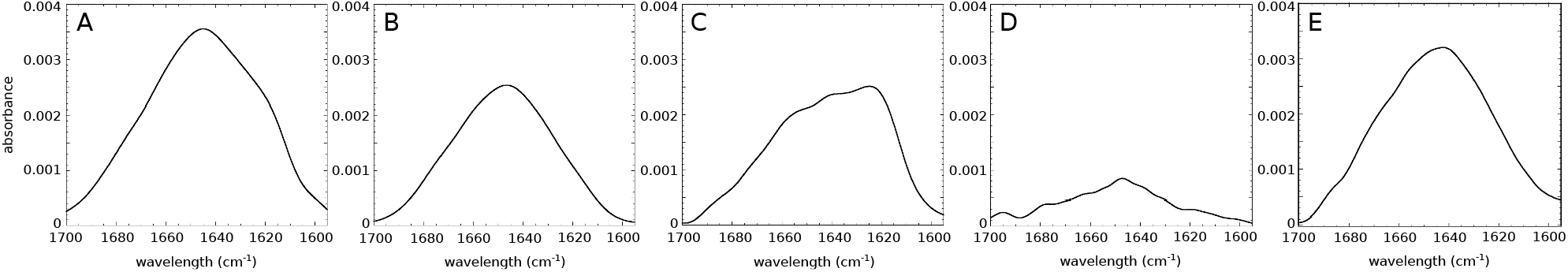
ATR-FTIR spectra for the amide I region of (A) hIAPP and the mutated peptides (B) H18R-IAPP, (C) H18K-IAPP, (D) H18E-IAPP, and (E) H18A-IAPP interacting with a lipid bilayer composed of DOPC/DOPS (7:3).

### 3.2 Influence of the residue 18 in IAPP conformational rearrangement

To evaluate the changes in secondary structure of the IAPP peptides at the DOPC/DOPS membrane interface, we collected ATR-FTIR spectra for 2 h in intervals of 30 min. Figure 2A shows that the maximum of the amide I band of hIAPP undergoes a pronounced shift from 1643 cm^−1^ to 1624 cm^−1^. This shift corresponds to a structural transition from an unstructured conformation to a structured one with antiparallel *β*-sheets, indicating the start of the peptide aggregation process. The maximum at 1624 cm^−1^ is reached at 120 min. However, a shoulder at around 1650 cm^−1^ remains, suggesting that not all of the amino acids are involved in the intermolecular *β*-sheet formation. The secondary structure content of the membrane-bound hIAPP at different incubation times (from 0 to 120 min) resulting from the analysis of the amide I band shape and curve fitting is given in Figure 2B. The bar chart clearly indicates that the *β*-sheet content increased from 31% to 50%, while the random coil content decreased from 37% to 26%, meaning that hIAPP started to aggregate, in agreement with previous studies Mishra et al. (2008). Nonetheless, some of the residues remained in an *α*-helical conformation, as this contribution dropped to ony about 15%, starting from 18% at time zero.

**Figure 2.**
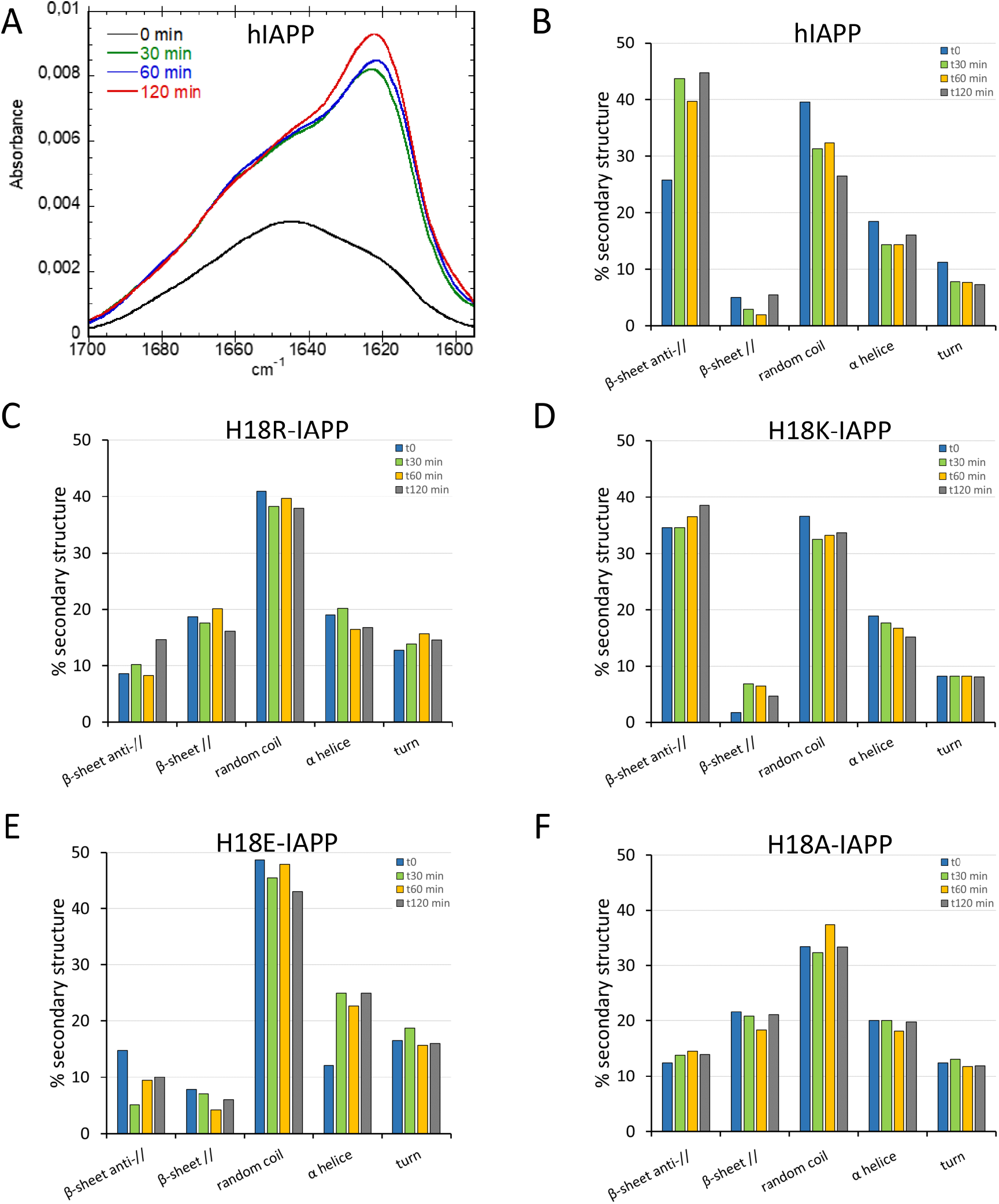
(A) Time-evolution (from 0 min, black, to 120 min, red) of the ATR-FTIR spectra of hIAPP. Secondary structure analysis for (B) hIAPP and the mutated peptides (C) H18R-IAPP, (D) H18K-IAPP, (E) H18E-IAPP, and (F) H18A-IAPP. The bars show the averaged content of secondary structures including antiparallel and parallel *β*-sheets, random coil, *α*-helices, and turns.

The same kind of experiments were performed for the mutated peptides. While they all undergo structural rearrangements, different behaviors are observed. As for hIAPP, the antiparallel *β*-sheet content of H18K-IAPP increases over time, thereby reducing the amount of random coil conformations, which suggests a self-assembly of the peptide (Figure 2D). It should be noted that already the structure of H18K-IAPP at the beginning of the experiment contains considerable amounts of antiparallel *β*-sheet, indicating that the aggregation of this peptide is immediate. Also in the cases of H18R-IAPP and H18A-IAPP there are *β*-sheets present at *t* = 0, yet they include both parallel and antiparallel arrangements, suggesting the presence of two structural populations (Figure 2C,F). These two populations are largely stable over time; only for H18R-IAPP some increase in antiparallel *β*-sheet content is observed at *t* = 120 min. In the case of H18E-IAPP, on the other hand, the amount of both parallel and antiparallel *β*-sheet decreases, whereas *α*-helical structures are increasingly formed (Figures 2E), reaching helical contents of more than 20%. This suggests that the DOPC/DOPS membrane promotes an *α*-helical conformation in membrane-bound H18E-IAPP. It should be mentioned that also in the case of H18A-IAPP the initial *α*-helix that formed remained stable, with population values of about 20%, whereas in H18K-IAPP and H18R-IAPP the helical content decreased somewhat to about 15%, which is similar as for hIAPP. In order to corroborate these results and validate the presence of one or two *β*-sheet populations, we then performed TEM in the presence of DOPC/DOPS membranes.

### 3.3 Electron microscopy images validate the structural differences between the peptides

TEM was applied to assess the presence of amyloid fibrils and/or amorphous aggregates interacting with the membrane. In the case of hIAPP, fibrils were obtained that exhibit a classical and mature amyloid-fibril morphology with widths of 6–10 nm (Figure 3A). It seems reasonable to assign these fibrils to the antiparallel *β*-sheets structure observed in the ATR-FTIR experiments. The same result is found for H18K-IAPP, where long and twisted fibrils are observed by TEM, which mainly harbor antiparallel *β*-sheets as revealed by the ATR-FTIR spectrum. For H8R-IAPP and H18A-IAPP, two aggregate morphologies are present in the TEM images, one corresponding to short fibrils and the other one being small amorphous aggregates (indicated by yellow arrows in Figure 3B,E). These results correlate with the ATR-FTIR experiments of both peptide variants that display two *β*-sheet populations: parallel and antiparallel *β*-sheets. Based on the observation that in the cases of hIAPP and H18K-IAPP the fibrils are correlated with the appearance of antiparallel *β*-sheets, we assume that also for H18R-IAPP and H18A-IAPP the antiparallel *β*-sheets give rise to fibrils, while the parallel *β*-sheets are most likely present in the amorphous aggregates. This suggests that amorphous aggregates and fibrils can not only be distinguished from each other, but they also arise from different secondary and tertiary structures. For H18E-IAPP, no fibrils were detected in the TEM images, only small aggregates occurred. The *β*-sheet content (both parallel and antiparallel) was also rather low; instead, the amount of random coil is rather high, suggesting that the amorphous H18E-IAPP aggregates are mainly unstructured. The curremt findings correlate with previous results that the substitution of the histidine 18 by an arginine, an alanine, or a glutamate stabilizes the oligomeric species and slows down the fibril formation Hoffmann et al. (2018). To gain more insights into the impact of residue 18 on the initial structure of the peptides and on the membrane interactions of IAPP and resulting structural changes, we performed all-atom MD simulations.

**Figure 3.**
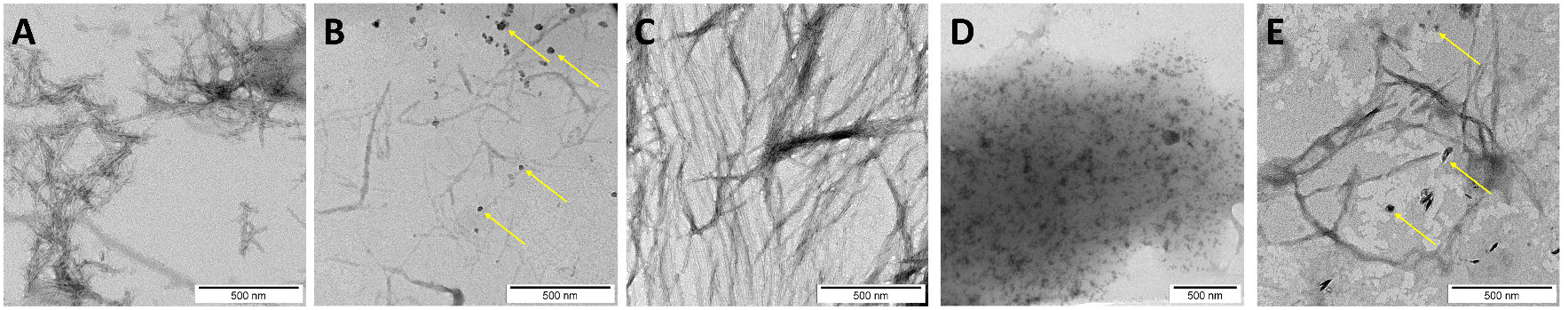
TEM image of (A) native hIAPP and the mutated peptides: (B) H18R-IAPP, (C) H18K-IAPP, (D) H18E-IAPP, and (E) H18A-IAPP incubated with DOPC/DOPS liposomes. The yellow arrows indicate the amorphous aggregates found for H18R-IAPP and H18A-IAPP. Scale bars represent 500 nm.

### 3.4 Atomic insight into the initial peptide structures at the membrane and their interactions from MD simulations

We performed 3 × 1 *µ*s MD simulations of hIAPP (with neutral H18 and positively charged H18, denoted as H18+) and its mutants H18A, H18E, H18R and H18K in the presence of a DOPC/DOPS (7:3) lipid bilayer.

#### 3.4.1 Membrane adsorption

To follow the association of the peptide with the membrane, we calculated the average distance between the center of mass of each residue and the average position along the bilayer normal of the phosphorus atoms of DOPC, which was used as a reference and therefore set to zero (Figure 4). It can be seen that the peptides interact differently with the membrane. Similar distance profiles are observed for hIAPP, hIAPP(H18+), H18R-IAPP and H18K-IAPP, while those of H18E-IAPP and H18A-IAPP are similar with each other yet differ from the other four. The smallest distances are witnessed for H18R-IAPP and H18K-IAPP, followed by hIAPP(H18+), which indicates that a positive charge at position 18 is key for its interaction with the membrane. The peptides generally approach the membrane with their N-terminus, with close contacts being formed between region K1–R19 and the lipids, while residues S20–Y37 are further away from the membrane. However, this does not apply to H18K-IAPP and H18R-IAPP, where almost all residues are within ≈1.0 nm of the membrane surface. In particular in the latter case, also the C-terminus is close to the membrane, indicating a parallel alignment of the peptide to the membrane surface, which is not as strongly visible for the other peptides. The profiles of the distance plots are characterized by a zigzag pattern, which suggests that the peptides adopt a helical structure on the membrane, that especially involves the first half of the peptides.

**Figure 4.**
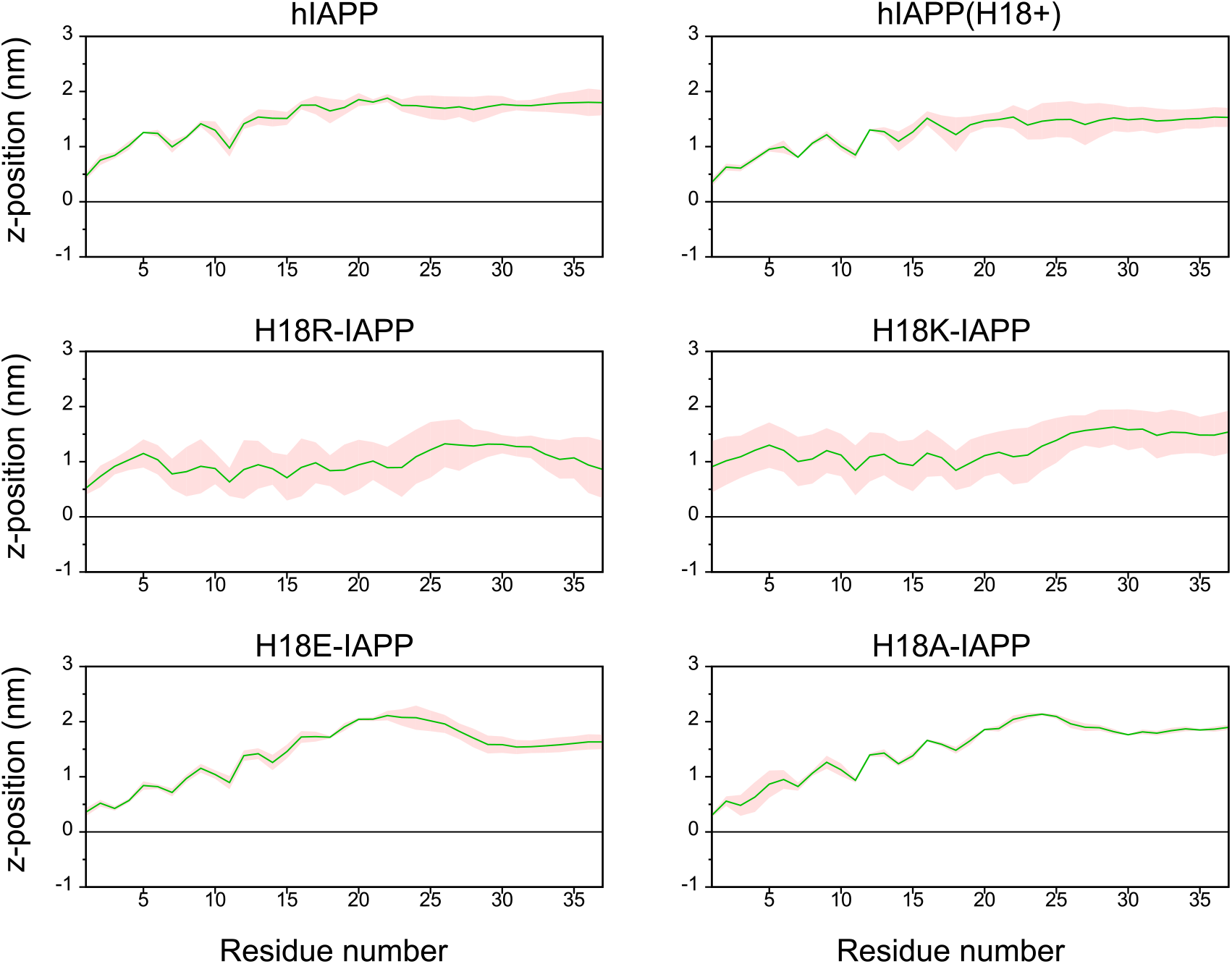
The average distance (and standard error, shown as shaded area) between each IAPP residue and the DOPC/DOPS membrane surface, which is defined by the average *z*-position of the P atoms of DOPC (shown as black line).

Figure 5 shows representative snapshots for the membrane association of IAPP, which confirms that the peptides tend to adopt a helical conformation in the N-terminal half. However, the wild-type peptides hIAPP and hIAPP(H18+) also involve a *β*-sheet in the C-terminal region (S20–G33), which was not adopted by the other peptides. In agreement to the distance plot one can see that hIAPP(H18+) inserts more deeply into the membrane than hIAPP, while hIAPP is only on, but not in the membrane. Nonetheless, in both cases the *β*-sheet interacts with the membrane, suggesting it to play a role in the subsequent aggregation when several peptides are membrane-adsorbed. These structures could even represent the *α*-to-*β* intermediate that was suggested to exist along the amyloid aggregation pathway of hIAPP, especially when this aggregation is assisted by the presence of lipid membranes (Abedini and Raleigh, 2009; Ling et al., 2009). Peptides H18R- and H18K-IAPP are seen to be immersed in the membrane. The helix, which reaches from T6 to G24 in both cases, lies below the lipid headgroup and is parallel to the membrane surface. In the case of H18R-IAPP, also the C-terminal residues are close to the headgroups, whereas the C-terminus of H18K-IAPP points away from the membrane surface, which explains the slight difference in their distance profiles shown in Figure 4. In the case of H18E-IAPP, the helix is least developed and all residues that are not part of the helix point away from the membrane. With the H18A mutation, on the other hand, a helix is formed, which however is not membrane-adsorbed. Only a few residues from the N- and C-terminal region make contact with the membrane, whereas the helix is several Angstrom above the membrane surface. The observation of a well-developed helix for H18A-IAPP is in line with the experimental findings and derives from the helix-promoting alanine introduced into the sequence. There are furter findings from the simulations that agree with the experimental results in Table 1. For instance, both simulations and experiments found that the *α*-helical content is smallest and that of random coil is largest for H18E-IAPP.

**Figure 5.**
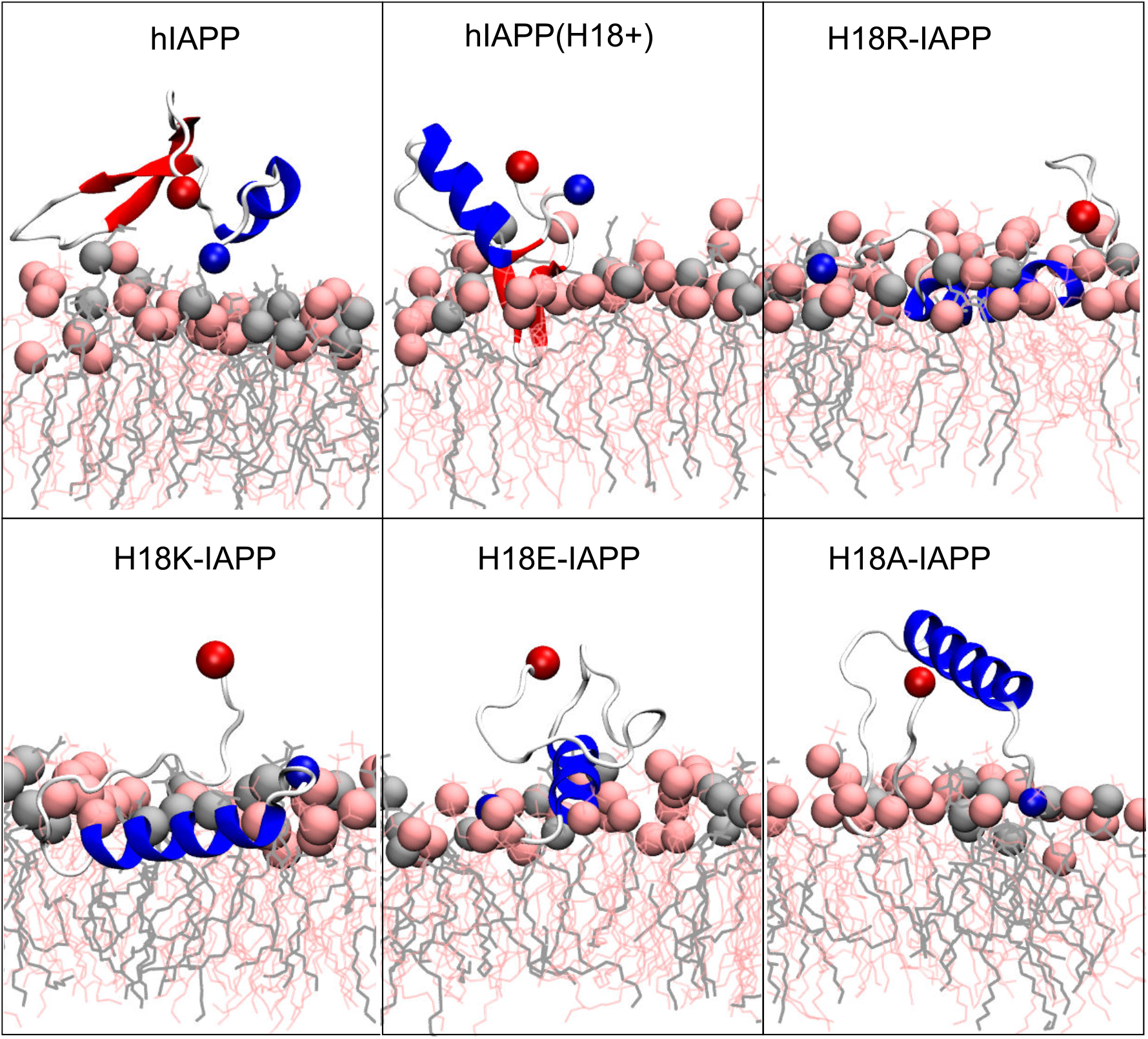
Representative IAPP structures interacting with the DOPC/DOPS membrane. The peptide is shown as cartoon (with helix, *β*-sheet and coil being shown in blue, red and white, respectively), with their N- and C-termini being indicated as blue and red spheres, respectively. DOPC and DOPS lipids are shown as pink and gray sticks, respectively, with their P atoms indicated by spheres of the corresponding color.

#### 3.4.2 Secondary structure

To quantify the effect of the peptide mutation and membrane adsorption on the peptide secondary structure, we determined the propensity of each peptide to adopt a helical conformation, to be part of a *β*-sheet (*β*-strand or *β*-bridge), or be in a turn or bend conformation. Figure 6 shows that random coil and *α*-helices are the dominating structures, with probabilities between 30 and 40%, or even above. Turn conformations are populated with a probability of 20–25%, while the *β*-sheet content is *<* 1%, apart for hIAPP where it is ≈ 5%. All of the mutants have a higher amount of helix than hIAPP. For H18K-IAPP it even reaches 45%, which correlates with its close interaction with the membrane. However, while for H18R-IAPP the interaction with the membrane is similar, the increase in helix is not as pronounced (35%). The second highest amount in helix is observed for H18A-IAPP (38%), which agrees to the increase in helical propensity seen for this mutant in experiment (Table 1) and results from the general helix-promoting characteristic of alanine. However, it should be mentioned that, despite all efforts, the comparability between simulation and experiment is limited. In simulation we only model the very first peptide–membrane interaction of the IAPP monomer, whereas in the ATR-FTIR experiments longer time scales and probably also small IAPP oligomers, in addition to monomers, were measured. This explains the generally low amount of *β*-sheet that is present in the simulated systems, as this is expected to increase upon IAPP aggregation. Only for hIAPP an average *β*-sheet content of 5% is observed that results from an intrapeptide *β*-hairpin that formed towards the end of the simulation. For hIAPP(H18+) it formed even later, therefore the average *β*-sheet content is lower even though for this system a *β*-hairpin is clearly visible in Figure 5.

**Figure 6.**
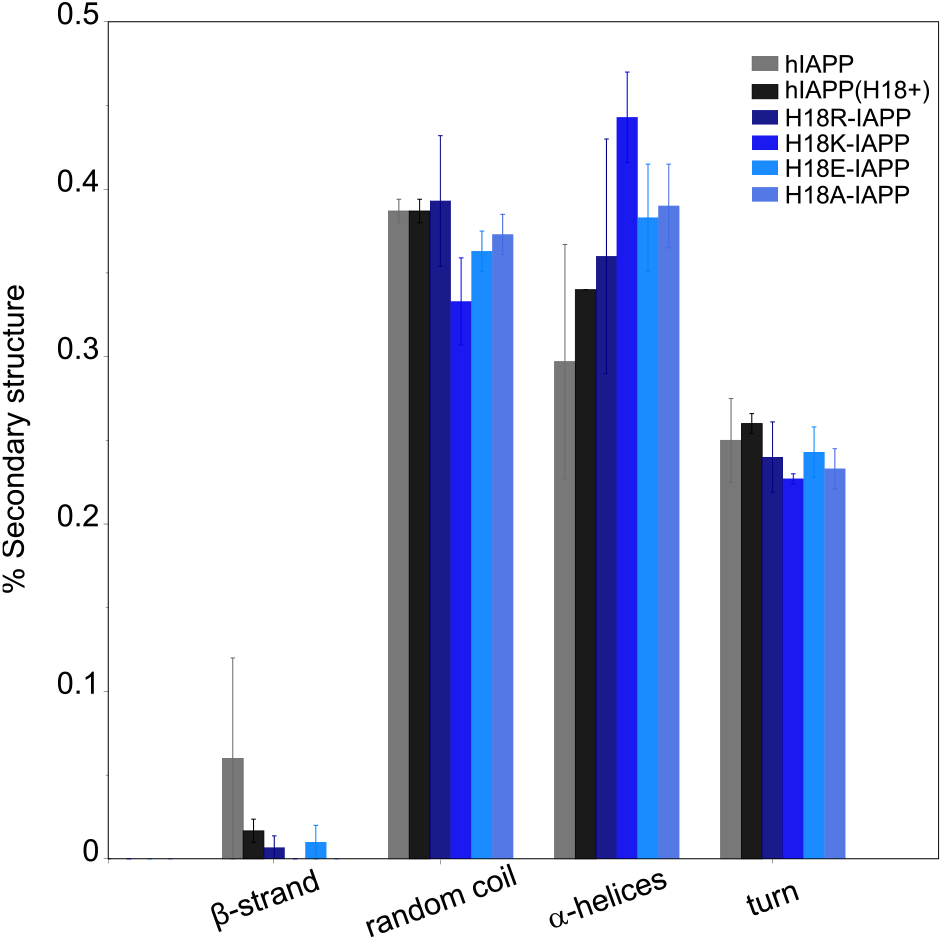
The average secondary structure content (and standard error) in IAPP in the presence of a DOPC/DOPS membrane as obtained from MD simulations.

#### 3.4.3 Peptide–membrane interactions

To rationalize the driving force for IAPP to interact with the DOPC/DOPS lipid bilayer, the interaction energy of each peptide residue with each component of the lipid bilayer was calculated and partitioned into its electrostatic (*E*_Coul_) and Lennard-Jones (*E*_LJ_) contributions (Supplemetary Figure S1). The results show that the major driving force for the peptide–membrane association are electrostatic attractions, especially between the negatively charged DOPS lipids and the positively charged residues K1 and R11. These interactions occur in all cases and explain why IAPP approaches the membrane always via its N-terminus. This observation agrees with those from previous MD studies that highlighted the importance of anionic lipids like POPG (1-palmitoyl-2-oleoyl-sn-glycero-3-phosphoglycerol), POPE (1-palmitoyl-2-oleoyl-sn-glycero-3-phosphoethanolamine), or DOPS in driving hIAPP–membrane interactions (Dignon et al., 2017b; Mei et al., 2020; Zhang et al., 2012). Supplemetary Figure S2 reveals that a positive charge at position 18 generally increases the tendency of the peptide to interact with the membrane. Almost all residues of the three peptides hIAPP(H18+), H18R- and H18K-IAPP form contacts with the membrane, whereas these contacts are mainly limited to K1–R11 in the other three cases. In the cases of H18R- and H18K-IAPP, the interaction between the positive charge of residue 18 and DOPS particularly enhances the association of the peptide with the membrane, which explains their deeper insertion into the membrane. This suggests that the size and/or flexibility of the side chain is important too. The electrostatic interactions partly involve H-bond formation in the region K1–R11, which extends to the C-terminal residues for hIAPP(H18+), H18R- and H18K-IAPP (Supplemetary Figure S3). Again, the positively charged residues are most involved in H-bond formation. The propensity of residue 18 to form an H-bond with DOPS or DOPC is particularly pronounced for H18R-IAPP, which is accompanied by further H-bonds between C-terminal residues and especially DOPC. Interestingly, in the experiments this peptide appeared to form fewer fifbrils and more amorphous aggregates compared to hIAPP and H18K-IAPP. Its tendency to form H-bonds with the lipids may explain why H18R-IAPP has a reduced propensity to form fibrils, which requires H-bonds to be formed between the peptides in order to enable *β*-sheet formation.

#### 3.4.4 Membrane insertion pathways

All-atom MD simulations allow to unravel the steps leading to the different peptide–membrane interactions in detail. An important aspect here is the high amount of hydrophobic residues present in IAPP, which give rise to an amphipathic helix when residues Q10 to L27 adopt an *α*-helix (Figure 7A). When such a helix binds to a membrane, it orients itself parallel to the membrane surface, with the hydrophobic side of this helix inserting into the hydrophobic core of the membrane and the hydrophilic residues of the other side interacting with the lipid headgroups or the aqueous solvent. This situation is visible for H18R-IAPP (Figure 7B). However, as the helix formed in this peptide only extends to S19, residues F23, I26 and L27 are not inserted into the membrane. Figure 7B further shows that the initial binding to the membrane is clearly driven by electrostatic interactions between the N-terminus and K1, which is followed by membrane insertion of the hydrophobic side of the amphipathic helix. This binding pattern is stabilized by interactions between R18 and the lipid headgroups, which is facilitated due to the length and flexibility of this side chain. For H18K-IAPP, the situation is similar, whereas in the case of hIAPP(H18+) the side chain is too short to enable strong interactions with the lipid headgroups. Figure 7C shows that this residue tends to be oriented toward the solvent. The interaction of hIAPP(H18+) with the membrane is dominated by K1, but the hydrophobic residues of the C-terminal side (F23, I26, L27) can also insert into the membrane, yet without forming a helix. Alternatively, these three residues can form a hydrophobic cluster, which can give rise to a *β*-hairpin as seen for both hIAPP and hIAPP(H18+) (Figure5).

**Figure 7.**
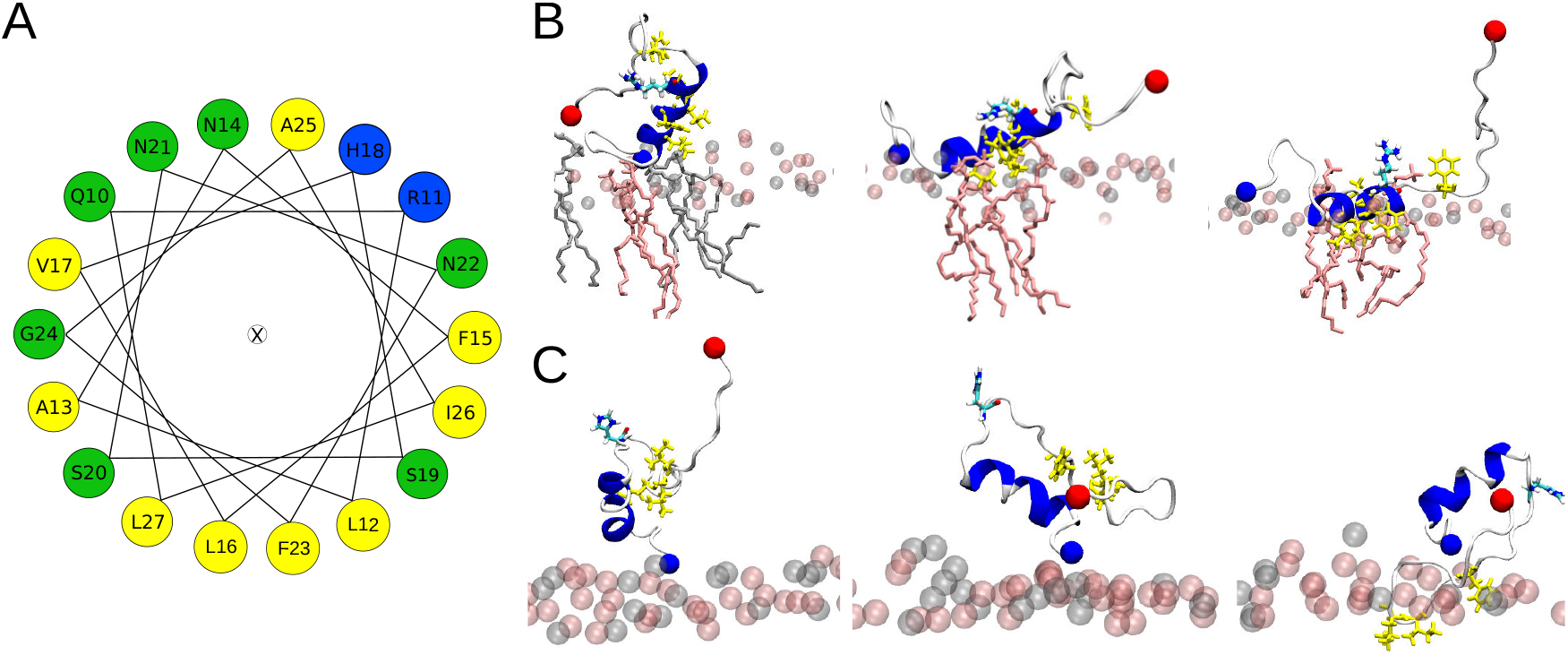
(A) Helical wheel of residues Q10 to L27 of hIAPP. Hydrophobic residues are highlighted by yellow, positively charged ones by blue, and polar residues by green. The orientation of the wheel was chosen such that the residues that insert into the membrane are located at the bottom. (B) This situation can be seen for IAPP-H18R. The initial membrane association is driven by electrostatic interactions between K1 and the lipid headgroups (left). Next, the hydrophobic residues (side chains shown in yellow) start inserting (middle) until their side chains are within the hydrophobic membrane core (right). This interaction is further stabilized by H-bonds between R18 and the lipid headgroups. (C) In the case of hIAPP(H18+) the side chain is too short for stable interactions between this residue and the membrane. Instead, this interaction is dominated by K1 (left and middle) or the hydrophobic residues F23, I26 and L27, while being in a non-helical state, insert into the hydrohpobic membrane core (right). The lipid headgroups are shown as spheres and in (B) the lipid heads close to the peptide are shown as sticks.

### 3.5 Effect of the peptides on the membrane properties

With polarized ATR-FTIR spectroscopy, not only the secondary structure of the peptides can be probed, also the effect of the peptides on the organization of the lipidic membrane can be determined. This is possible by measuring the position of the bands corresponding to antisymmetric and symmetric stretching modes of the methylene groups of the lipid tails, *ν*_as_(CH_2_) and *ν*_s_(CH_2_) in the absence and in the presence of the peptides as well as the dichroic ratio (*R*_ATR_) of the *ν*_s_(CH_2_) bands (Table 2) Goormaghtigh et al. (1999). The wavenumbers of these bands are known to be sensitive to changes in the configuration of the acyl chains, in chain mobility, and packing. For the bilayer alone, *ν*_s_(CH_2_) and *ν*_as_(CH_2_) are 2854 and 2945 cm^−1^, respectively, and the value of *R*_ATR_ is 1.28, which is characteristic for fluid and packed acyl chains. The addition of hIAPP to the bilayer does not significantly change the wavenumbers, while there is a slight increase in *R*_ATR_ for the *ν*_s_(CH_2_) bands, which indicates a minor increase in disorder in the lipid chains. In contrast, in the presence of the mutated peptides, the wavenumbers are not modified, suggesting that these peptides do not or hardly affect the organization of the lipid bilayers. Thus, our results show that the mutated peptides do not alter the membrane properties during the first peptide–membrane interaction events, while hIAPP slightly increases the disorder in the membrane resulting from initial peptide insertions into the membrane.

**Table 2.** Wavenumbers and dichroic ratio for the methylene groups of the lipid chains in the absence and presence of IAPP peptides.

| | $\nu_{\text{as}}(\text{CH}_2)$ ($\text{cm}^{-1}$) | $\nu_{\text{s}}(\text{CH}_2)$ ($\text{cm}^{-1}$) | $R_{\text{ATR}}(\nu_{\text{s}}(\text{CH}_2))$ |
| --- | --- | --- | --- |
| supported lipid bilayer | 2924 | 2854 | $1.28 \pm 0.07$ |
| + hIAPP | 2919 | 2851 | $1.34 \pm 0.07$ |
| + H18R-IAPP | 2927 | 2855 | $1.29 \pm 0.07$ |
| + H18K-IAPP | 2927 | 2855 | $1.27 \pm 0.07$ |
| + H18E-IAPP | 2925 | 2854 | $1.26 \pm 0.07$ |
| + H18A-IAPP | 2926 | 2855 | $1.20 \pm 0.07$ |

This conclusion is supported by the analysis of the membrane bilayer properties from the MD simulations. From the mass density profiles of the DOPC and DOPS headgroups along the membrane *z*-axis we determined the average bilayer thickness as ≈4 nm. In order to assess whether the peptides affect the membrane properties, we analyzed the bilayer thickness around the membrane-associated peptides (Supplemetary Figure S4). In these areas, reductions in the bilayer thickness of up to ≈0.1 nm are detected. However, hIAPP and H18A-IAPP have only small to no effects on the bilayer thickness, suggesting that a charged amino acid at position 18 plays a role in causing membrane perturbations, especially when it enables membrane insertion. This is best seen for H18R- and H18K-IAPP that triggered the largest changes in membrane thickness, which are the same two peptides that inserted into the membrane during the simulations. For further characterization of the membrane properties, we calculated the order parameter of the C–H bonds in the lipid acyl chains (denoted as *S*_CH_) for both DOPC and DOPS lipids. Here, we distinguished between lipids that are in the vicinity of the peptides (i.e., within 0.5 nm) and all other lipids, to observe whether the peptides can cause lipid disorder (Supplemetary Figure S5). Similar *S*_CH_ profiles along the acyl chains (characterized by carbon number) are observed for DOPC and DOPS, with the order parameters of the latter being slightly higher. A strong drop in order is present at the double bonds positioned at carbon atom 10 of both palmitoyl and oleoyl chains of either lipid type. Most importantly, no notable change in lipid order due to the presence of any of the peptides is observed. This suggests that changes to the lipid thickness resulted only from the interactions between the peptides and the lipid headgroups, while the acyl chains are not affected as none of the peptides did insert deeply into the membrane core, maximally just below the headgroup region in the cases of H18R- and H18K-IAPP.

Apart from hIAPP this agrees to the observations from the experiments, as also there the lipid tail packing was not affected by the peptides, suggesting that also in the experiments the peptides did not penetrate into the membrane core. Only in the case of hIAPP some minor changes in the acyl packing were recorded, indicating that in the experiments this peptide was able to notably reach beyond the headgroup region. Interestingly, this is the same peptide that formed the largest amounts of (antiparallel) *β*-sheets, suggesting that *β*-sheet formation and membrane insertion take place concurrently.

## 4 CONCLUSIONS

In the present study, ATR-FTIR and TEM experiments as well as all-atom MD simulations on the microsecond time scale have been performed to unravel the first structural changes of the islet amyloid polypeptide following its interaction with a lipid membrane. Moreover, the influence of residue 18 in this process was assessed by studying wild-type hIAPP and variants of it with mutations H18A, H18E, H18K, and H18R.

The secondary structure profiles from simulations suggest that initially the membrane-bound IAPP is mostly in *α*-helical and random coil conformations (≈30–40%). All mutants show higher amounts of helix than hIAPP, especially H18K-IAPP (≈45%) and H18A-IAPP (≈38%). While alanine is commonly known to be a helix-promoting amino acid, also lysine has a high helix-forming propensity. The notably lower amounts in *β*-sheet in the simulations compared to what is observed experimentally is due to the different length and time scales that are assessed. The simulations are limited to one peptide and one microsecond and, hence, cannot sample peptide aggregation, which however takes place to a certain extent in the experiments. The analysis of the ATR-FTIR spectra demonstrates that at the beginning of the experiments, the peptides are predominantly unstructured (35–50%) with some contributions from *α*-helical structures (12–20%) and *β*-sheets. During the course of 2 h of incubation, the ATR-FTIR spectra of hIAPP revealed an increase in the antiparallel *β*-sheet content and a reduction in the *α*-helical and random coil contents, which is in agreement with the TEM images that revealed fibrils with typical amyloid morphology. Similar data as for hIAPP were obtained for H18K-IAPP that also experienced an increase in anti-parallel *β*-sheet content and exhibited typical amyloid fibrils. A solid-state NMR study demonstrated that hIAPP(20–29) amyloid fibrils contain anti-parallel *β*-sheet structures Griffiths et al. (1995). However, recent cryo-EM structures of full-length IAPP fibrils showed that the *β*-sheets are in a parallel configuration Röder et al. (2020); Gallardo et al. (2020). A difference is that the solid-state NMR and cryo-EM structures were determined for fibrils grown in solution, while here we studied IAPP fibrils that formed on supported lipid bilayers. We thus conclude that the bilayers modulate the amyloid fibrillation of IAPP in a way that antiparallel *β*-sheet are favored.

Previous reports revealed that hIAPP and H18K-IAPP are toxic to *β*-cell lines Khemtemourian et al. (2017). Based on our current observations, we suggest that the toxic hIAPP and H18K-IAPP species are those that are structured with antiparallel *β*-sheets, as different behaviors are observed for the other peptides. In the case of hIAPP, the joint analysis of the experimental and simulation data suggests that the *β*-sheet aggregates even started to insert into the lipid bilayers, causing membrane disorder, which would explain their toxicity. For H18R-IAPP and H18A-IAPP, both the ATR-FTIR spectra and TEM images indicate the presence of two species, one of them being structured in antiparallel *β*-sheets and the other one involving parallel *β*-sheets. The TEM images further revealed two different supramolecular structures, thin and short fibrils as well as amorphous aggregates. We propose that the fibrils are composed of the antiparallel *β*-sheets, as seen for hIAPP and H18K-IAPP, while the amorphous aggregates contain parallel *β*-sheets. In all four peptides, the formation of *β*-sheets was accompanied by a reduction in random coil – this was especially the case for hIAPP – and *α*-helix. Nonetheless, the amount of helix remained at 15–20%, suggesting that the initial membrane-anchoring via a helix in the N-terminal half of the peptide, as revealed here by simulations, is very stable and resists the transformation into *β*-sheets. The *β*-sheet formation is likely to take place in the other residues, such as IAPP(20–29) as was previously revealed by solid-state NMR spectroscopy. H18E-IAPP is the only peptide for which no transitions into *β*-sheets were observed, neither in the ATR-FTIR spectra nor did fibrils occur in the TEM images. Instead, the amount of helix increased with time. This concurs with the previous finding that H18E-IAPP is the least toxic H18 mutated peptide Khemtemourian et al. (2017) and reinforces our hypothesis that cytotoxicity and the presence of antiparallel *β*-sheet structures are correlated in IAPP. For other amyloid proteins such correlation has already been demonstrated. Using a yeast amyloid from the HET-s prion domain of *Podospora anserine*, Cullin and coworkers showed that mutations within the HET-s prion domain give rise to antiparallel *β*-sheet structures and, at the same time, enhance the cytotoxicity Berthelot et al. (2011). Other toxic amyloid-forming proteins also adopt an antiparallel *β*-sheet conformation, such as the amyloid-*β* peptide involved in Alzheimer’s disease and *α*-synuclein related to Parkinson’s disease Celej et al. (2012); Cerf et al. (2009), suggesting that the antiparallel *β*-sheet is a signature of amyloid toxicity.

Earlier studies indicated that hIAPP and the mutated peptides are able to induce membrane permeability Hoffmann et al. (2018). Here, we tested for possible membrane disorder induced by the peptide using both ATR-FTIR and MD simulations. Consistent with previous MD studies, the peptides approache the membrane via their N-terminal residues (K1–R19). This interaction between peptide and membrane is mainly driven by electrostatic attractions between the positively charged residues K1 and R11 and the negatively charged lipid DOPS, which is strengthened when there is a third positive charge at position 18, as seen for H18K- and H18R-IAPP. However, hIAPP(H18+) did not interact more strongly with the membrane, suggesting that, in addition to the charge at residue 18, also the size and/or flexibility of the side chain plays a role in affecting peptide–membrane interactions. In the simulations, the helical regions of H18K- and H18R-IAPP were able to insert into the membrane, adopting a parallel orientation with respect to the membrane surface where the hydrophobic side chains entered the hydrophobic membrane core and the hydrophilic side chains point in the opposite direction towards the aqueous phase. This orientation is stabilized by the long and the flexible side chains of K1, R11, and K18 or R18. The ATR-FTIR results reflect that the mutated peptides H18R-, H18K-, H18E-, and H18A-IAPP do not alter the membrane properties during the initial peptide–membrane interactions, while hIAPP was able to slightly change the membrane properties. Since this is the peptide with the largest amount of *β*-sheets being formed, this suggests that *β*-sheets are needed for membrane disturbances. This conjecture is further supported by our MD data which revealed that the initial insertion of IAPP as a helix is only just below the headgroup region, which, apart from small effects on the membrane thickness around the peptide, does not change the lipid tail order. This agrees to the findings from the ATR-FTIR spectra. Hence, we conclude that apart from hIAPP, no deep insertions of the peptides into the membrane occurred in the current experiments. Various membrane damage mechanisms caused by hIAPP have been proposed and described in detail, which implicate the presence of large oligomers or fibrils and involve pore formation or lipid uptake Engel (2009). Our experimental and simulation results indicate that the initial IAPP aggregate species are not able to inflict such membrane destabilization.

In summary, the results of this study provide valuable molecular level insight into understanding of the initial IAPP–membrane interactions and demonstrate how mutations at residue 18 can affect this interaction and fibril formation of IAPP. We demonstrated that a single mutation of histidine 18 can yield vastly different results in terms aggregate morphology, membrane damage and resulting toxicity, highlighting once again the importance of this residue in amyloid formation by hIAPP.

## AUTHOR CONTRIBUTIONS

L.K. designed the study. L.K performed the TEM and analyzed the images. L.K. and B.D. performed the ATR-FTIR experiments; L.K., S.L and S.C. analyzed the ATR-FTIR spectra. H.F. and B.S. designed the computer simulations. H.F. performed the simulations and analyzed them together with B.S. L.K., H.F. and B.S. wrote the manuscript. All authors contributed to discussing the results and reviewing the manuscript.

## ACKNOWLEDGMENTS

Christophe Piesse (Institut de Biologie Paris Seine, Université Pierre et Marie Curie, France) is acknowledged for the peptides synthesis. H.F. and B.S. gratefully acknowledge the computing time granted through JARA on the supercomputer JURECA at Forschungszentrum Jülich (project number JICS6C).

## SUPPLEMENTAL DATA

The Supplementary Material for this article can be found online.

