## Supplementary Figures for "Structural dissection of the first events following membrane binding of the islet amyloid polypeptide"

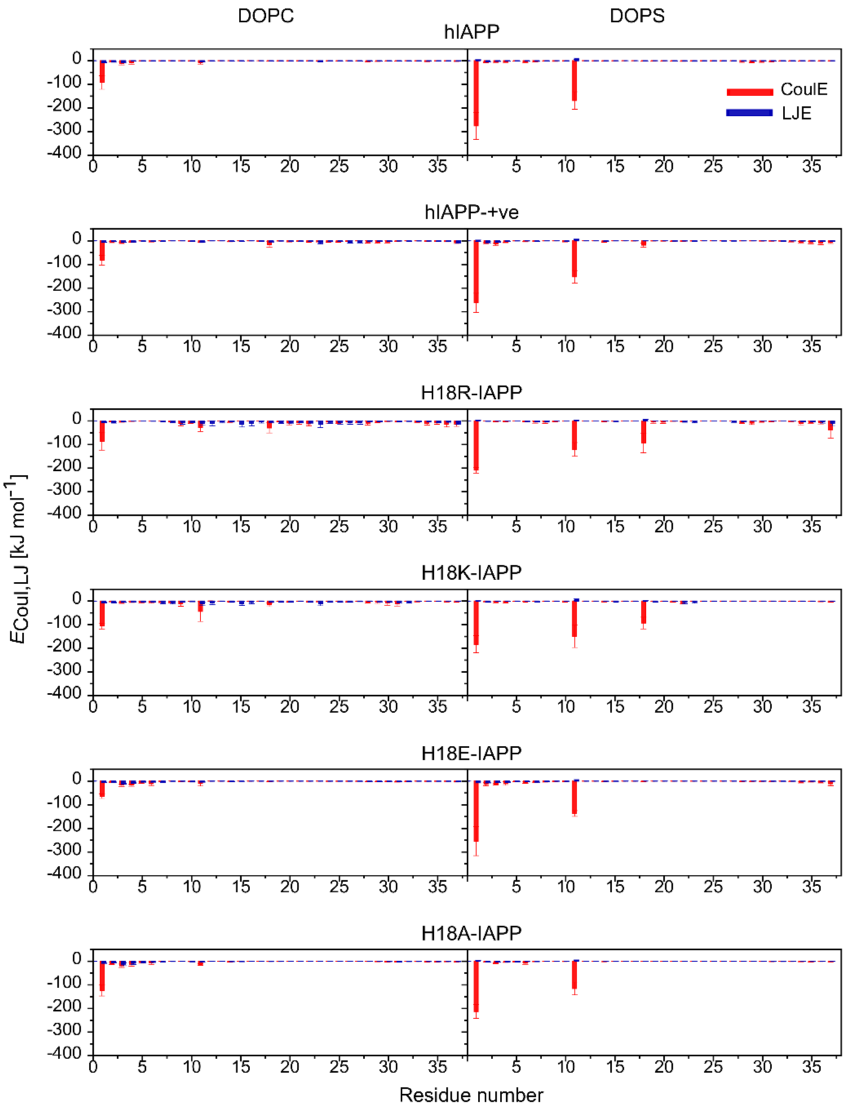


**Figure S1:** The average interaction energies (and standard error) of IAPP interacting with DOPC (left) and DOPS (right) lipids. Electrostatic and Lennard-Jones energies are shown in red and blue, respectively. Negative energies indicate attractive forces, positive energies correspond to repulsion.

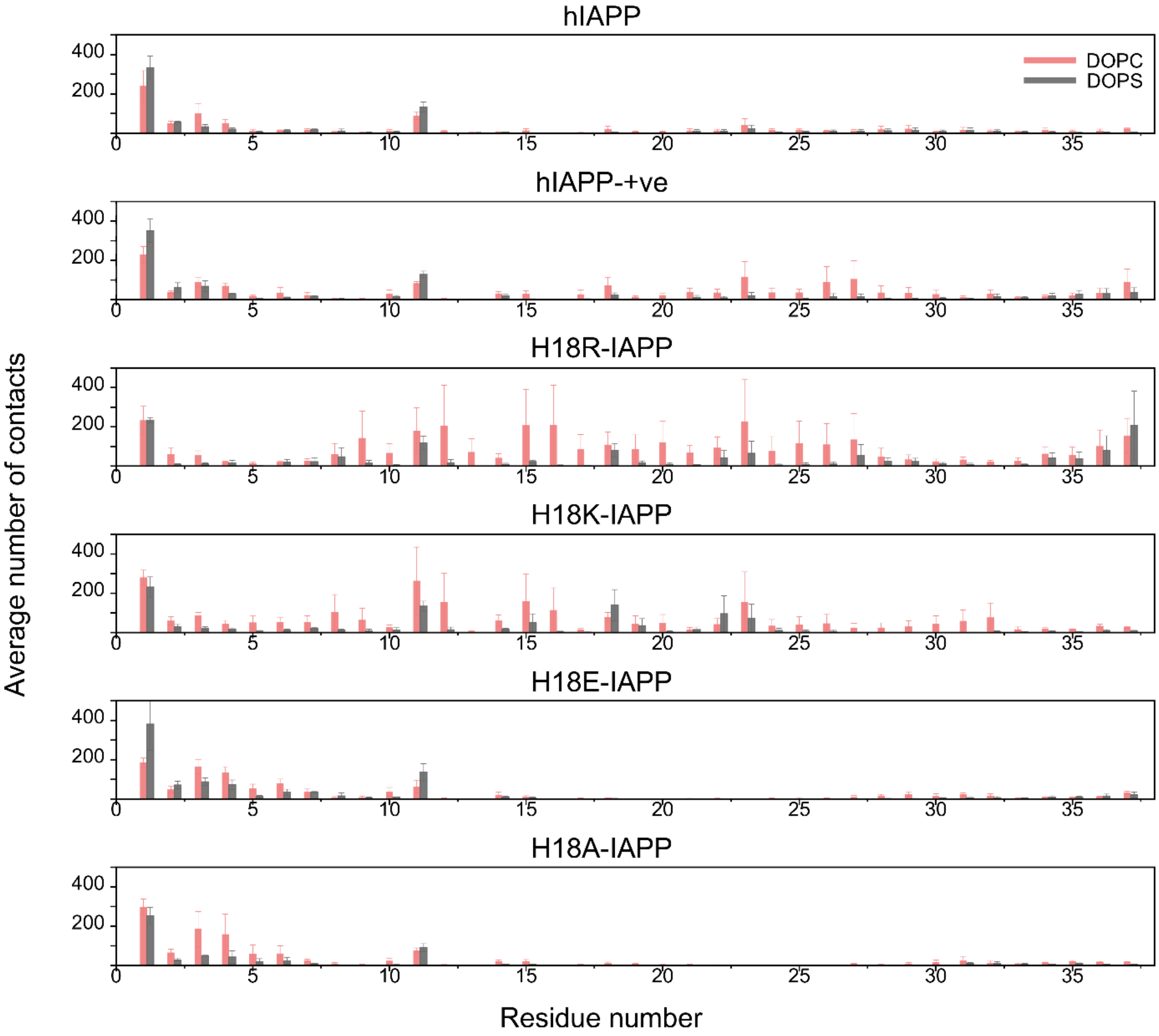


**Figure S2:** The average number of IAPP-lipid contacts (and standard error) for DOPC (pink) and DOPS (gray) lipids.

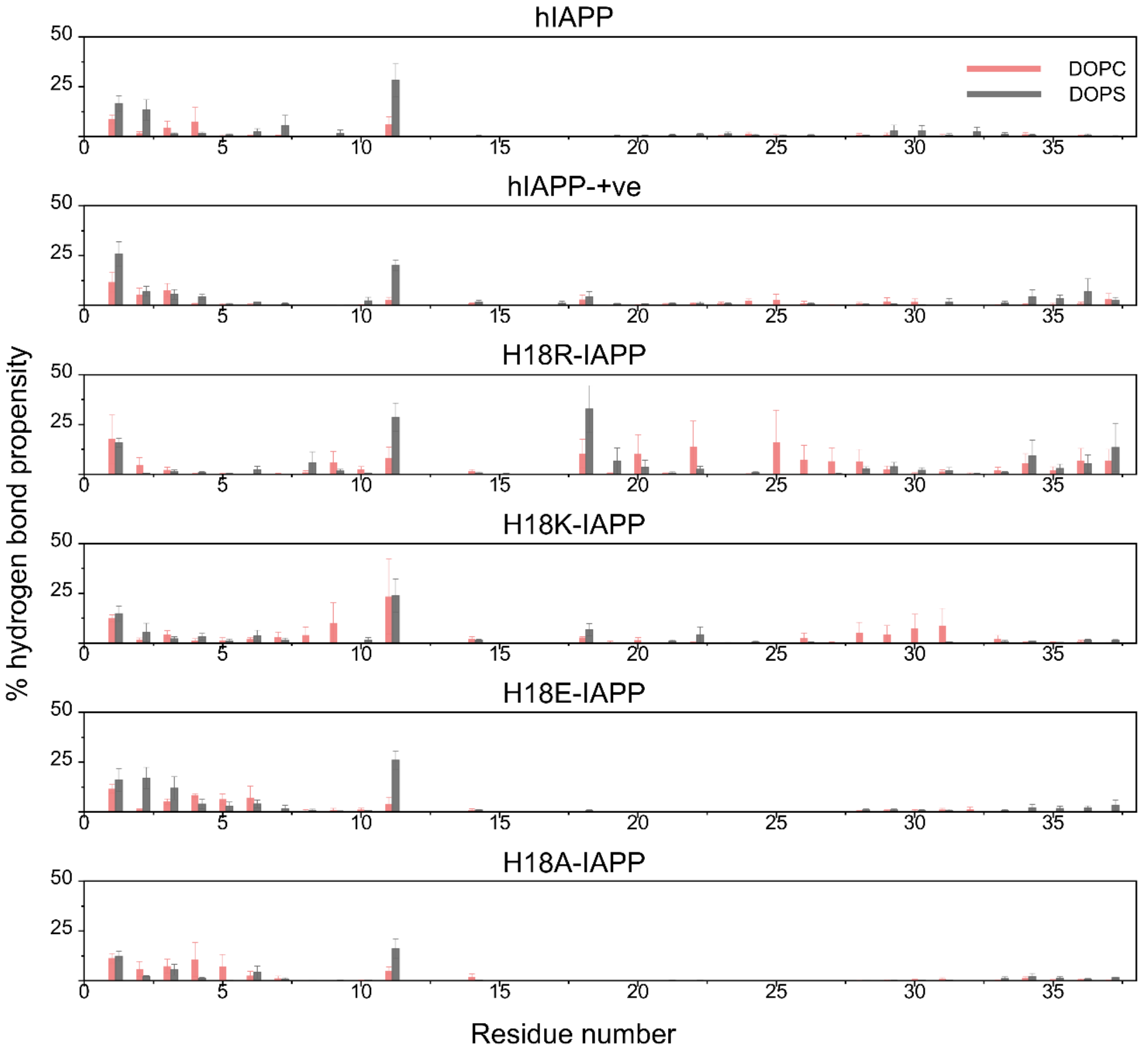


**Figure S3:** The average hydrogen bond propensity (and standard error) between IAPP and DOPC (pink) and DOPS (gray) lipids.

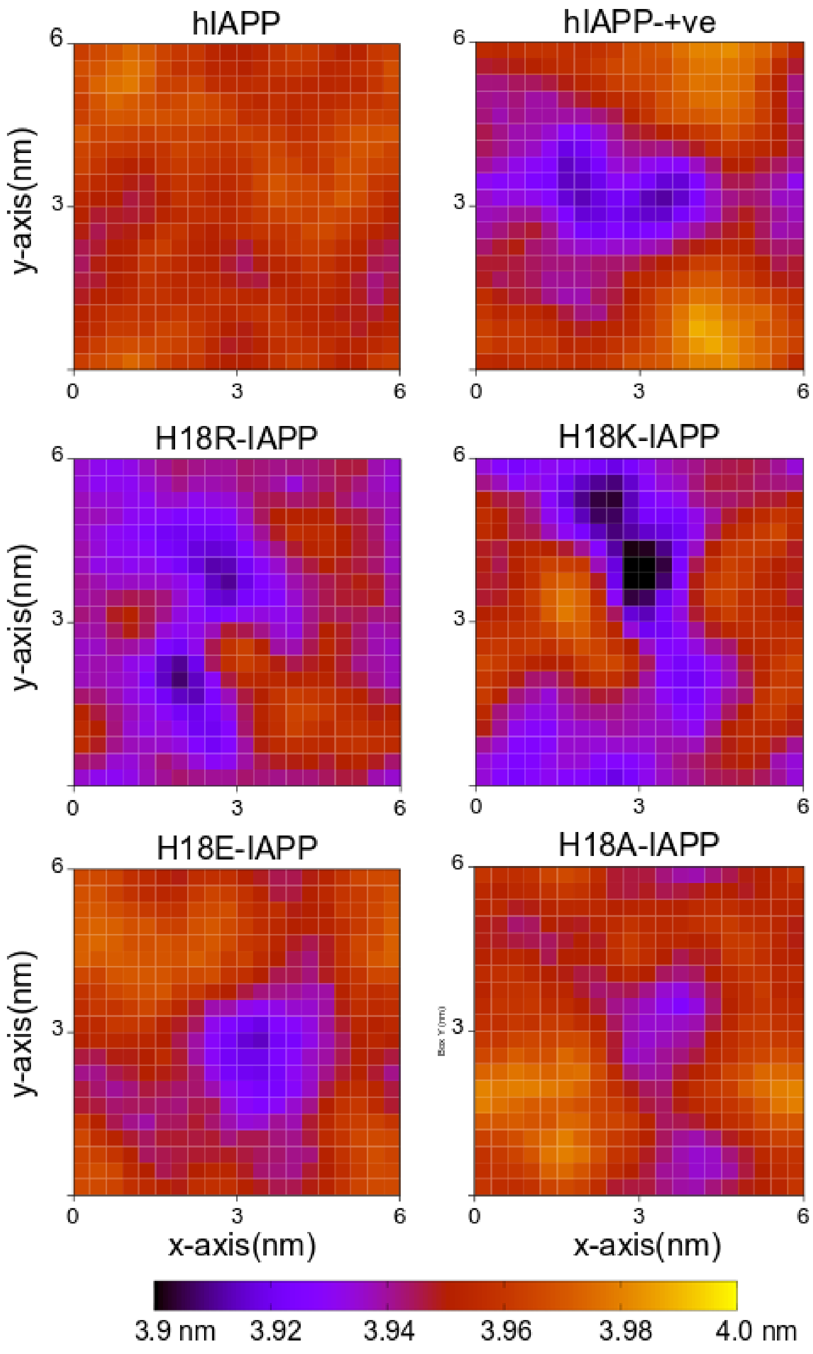


**Figure S4:** Average bilayer thickness calculated for the MD frames where the peptide is within 0.5 nm of the membrane. The *x*- and the y-axes represent the unit cell dimension in nm. The color bar shows the thickness range in nm.

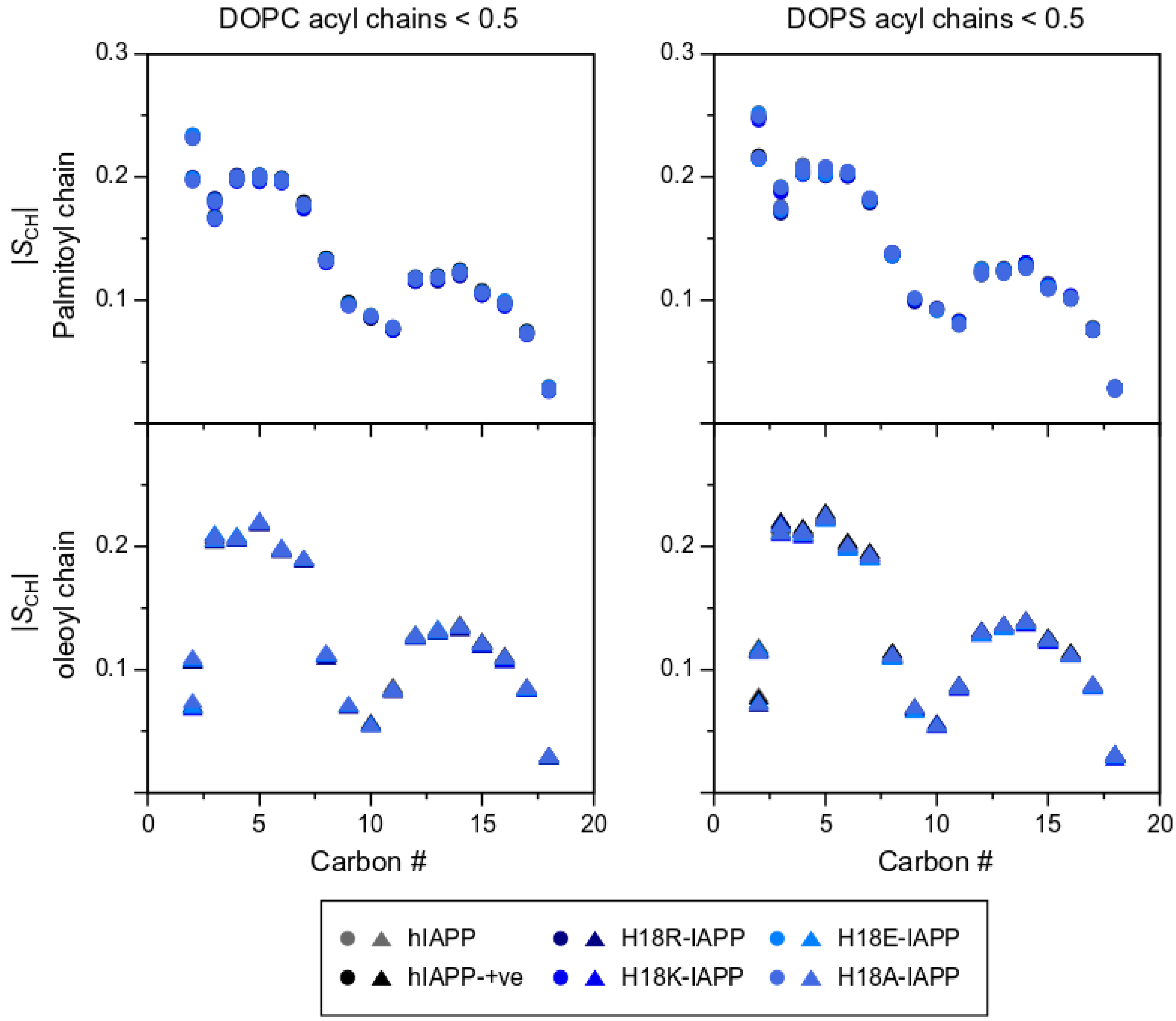


**Figure S5:** Average order parameters of the acyl chains (top: palmitoyol chains; bottom: olyeol chains) of DOPC (left) and DOPS (right) lipids.
